# The chromatin remodeler SMARCD3 regulates cell cycle progression and its expression predicts survival outcome in ER+ breast cancer

**DOI:** 10.1101/684217

**Authors:** Romain Tropée, Bárbara de la Peña Avalos, Madeline Gough, Cameron Snell, Pascal H.G. Duijf, Eloïse Dray

**Affiliations:** IHBI, Queensland University of Technology, Brisbane QLD Australia; Department of Biochemistry and Structural Biology, University of Texas Health Science Center at San Antonio, San Antonio, TX, USA; Cancer Pathology Research Group, Mater Research Institute – The University of Queensland, Translational Research Institute, 37 Kent St, Woolloongabba, QLD 4102; Mater Pathology, Mater Hospital Brisbane, Mater Misericordiae Limited, South Brisbane, QLD 4101; University of Queensland Diamantina Institute, The University of Queensland, Brisbane QLD Australia; Mays Cancer Center, UT Health San Antonio MD Anderson, San Antonio, TX, USA

## Abstract

Chromatin remodeling plays an essential role in regulating transcriptional networks and timing of gene expression. Chromatin remodelers such as SWItch/Sucrose Non-Fermentable (SWI/SNF) harbor many protein components, with the catalytic subunit providing ATPase activity to displace histones along or from the DNA molecules, and associated subunits ensuring tissue specificity and transcriptional or co-transcriptional activities. Mutations in several of the SWI/SNF subunits have been linked to cancer. Here, we describe how *SMARCD3*/Baf60c expression is associated with hormone positive (ER+) breast cancer. The level SMARCD3, as detected by immunohistochemistry in breast cancer patient samples, is correlated with differential long-term disease-free survival. In contrast, the expression level of *SMARCD1*/Baf60a and *SMARCD2*/Baf60b, which are mutually exclusive within the SWI/SNF complex and have a partially redundant function, lacks predictive value in breast cancer patient samples. Lower proliferation rates are observed in SMARCD3 depleted cells, which reflects a failure to fully progress through G2/M, and an increase in endoreplication. In the absence of SMARCD3, p21 accumulates in cells but does not halt the cell cycle, and DNA damage accumulates and remains unrepaired. Taken together, our data begin to explain why ER+ breast cancer patients with low SMARCD3 expressing tumors exhibit reduced survival rates compared to patients expressing normal or higher levels of SMARCD3. SMARCD3 might act as a tumor suppressor role through regulation of cell cycle checkpoints and could be a reliable and specific breast cancer prognostic biomarker.

**Significance:** Mutations in chromatin remodelers are a leading cause of cancer. Estrogen Receptor positive (ER+) breast cancers represent approximately 80% of all cases diagnosed. Although these tumors can be treated with hormone therapy, most breast cancer fatalities occur in ER+ breast cancer patients, due to metastasis. Low expression of SMARCD3 in ER+ cancer is associated with diminished survival rates. As such, SMARCD3 could be used as a predictive biomarker for survival. In addition, we have identified a role for SMARCD3 in the cell cycle, which could at least partially explain its protective role in breast cancer. While catalytic subunits are often viewed as the major components in chromatin remodeling function, we show here new evidence that mutations or silencing of SMARCD3 may also contribute to genomic instability and thus development of breast cancer.

## Introduction

The chromatin remodeler SWI/SNF harbors a large number of subunits (12 genes in yeast, 29 in human; see (1) for review; Fig. 1A), whose functions are conserved among eukaryotes (2). Early work identified SWI/SNF as a transcription activator through histones sliding and alteration of the nucleosome structure (3). Since then, it has been implicated in DNA replication, DNA repair, apoptosis, metabolism, and cell differentiation. Mutations and aberrant expression of SWI/SNF subunits are frequently found in cancer. Importantly, SWI/SNF mutations are linked to many cancer subtypes (4), hinting at subtle and tissue-specific regulation of the SWI/SNF subunits. Work done by a number of research laboratories has investigated the contribution of individual subunits in yeast (5, 6) and humans (7) in relation to the known biological functions of SWI/SNF, and interrogated the role of subunits in maintaining both the structural and functional integrity of the complex. These studies have provided insights into how mutations in subunits affect gene expression and favor cancer progression (5–7).

**Figure 1.**
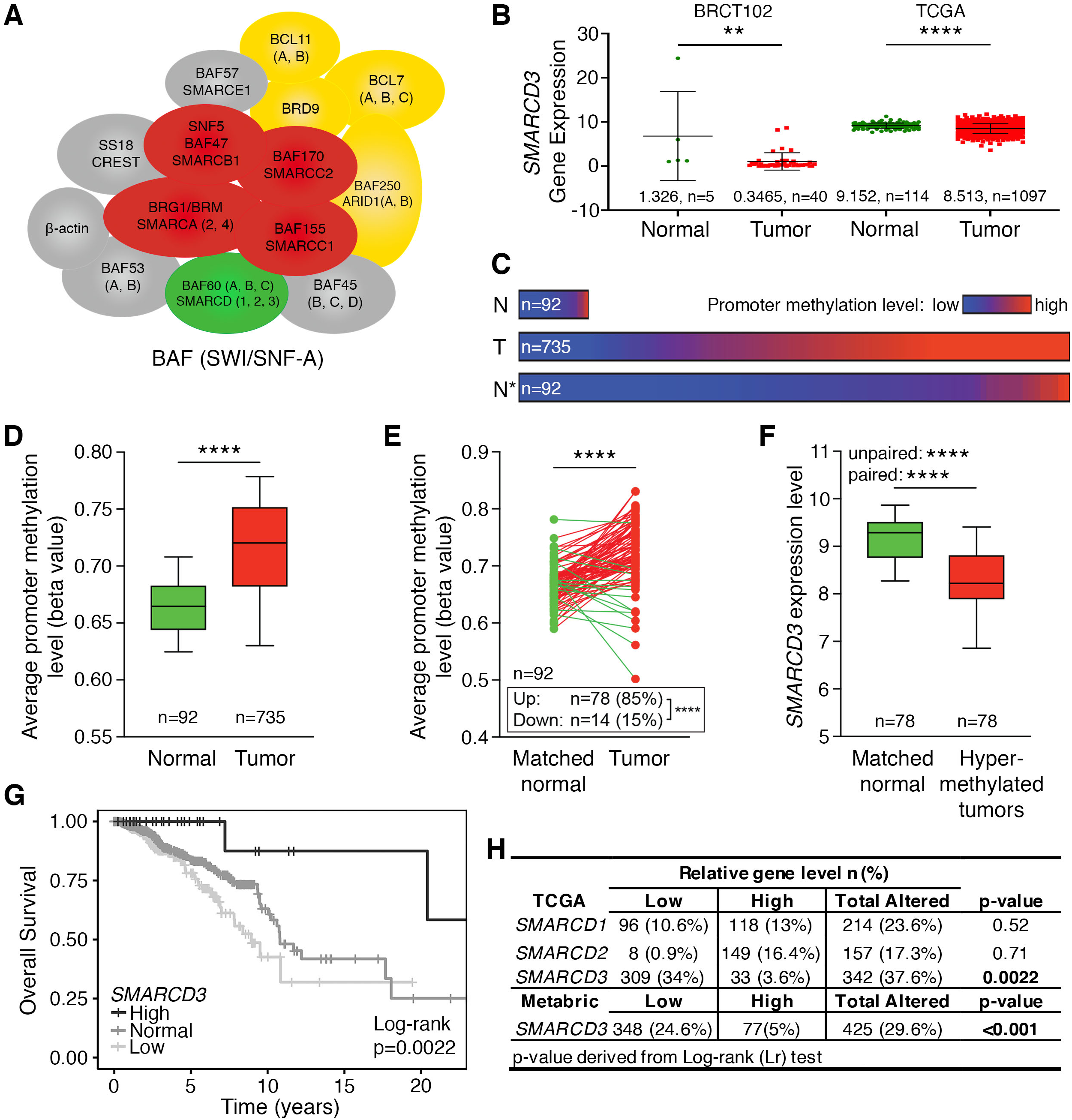
*SMARCD3* expression is modified in breast cancer. **(A)** Schematic of the BAF complex showing possible combinations of subunits (SMARCD1/2/3 in green). **(B)** RT-qPCR analysis of *SMARCD3* expression in BRCT102 relative to 18S and RNAseq analysis of *SMARCD3* in TCGA breast invasive carcinoma dataset (unpaired t-test, **\*\*** p < 0.01; **\*\*\*\*** p < 0.0001). **(C)** SMARCD3 promoter methylation levels are shown for all 92 normal (N) and all 735 tumor (T) samples (drawn to scale) in the TCGA Breast Invasive Carcinoma dataset. The degree of methylation is color-coded, as indicated in the key. To facilitate comparison of proportions between normal and tumor samples, normal samples are also shown off scale, stretched to same length as the tumor sample bar (N*). **(D**,**E)** The *SMARCD3* promoter is hypermethylated in TCGA breast cancer samples compared to normal samples. **(D)** Unpaired analysis (Mann Whitney U test). **(E)** paired analysis (Wilcoxon signed rank test). **(F)** Breast cancers with hypermethylated SMARCD3 promoters express significantly lower SMARCD3 mRNA than their respective matched normal samples, as per unpaired and paired tests (Mann Whitney U test and Wilcoxon signed rank test, respectively). **(G)** Kaplan Meier curves of overall survival in TCGA dataset, patient stratified using *SMARCD3* high, normal or low expression. Log-rank p-value testing for similarity between survival curve. **(H)** Frequency of patient with gene expression alteration in TCGA Breast Invasive Carcinoma and corresponding log-rank P val. testing for similarity between survival curve.

Much attention has focused on the association of SWI/SNF mutations or its epigenetic silencing in cancer. The catalytic subunits, BRG1 or BRM, are essential for enzymatic function and the structural integrity of SWI/SNF, and have been intensely studied. In contrast, tissue-specific subunits (BRG1 associated factors, or BAF) and non-catalytic subunits of SWI/SNF, e.g. SMARCD3, one of three paralogs of SMARCD (*SMARCD1/2/3*, or Baf60a/b/c respectively), remain poorly characterized. SMARCD1/2/3 are mutually exclusive, tissue-specific, and their function is only partially redundant as they cannot complement each other (8, 9). The complex containing SMARCD3 is found especially enriched in cardiomyocytes, where SMARCD3 is essential for the contractile function of cardiac cells (10). SMARCD3 is critical for cell differentiation *via* direct interactions with partners such as MYOD, thus coordinating cell differentiation cascades through transcriptional activation of genes, and through epigenetic reprogramming (11, 12). Recent studies have associated SMARCD3 levels and breast cancer risks: first, the promoter region of *SMARCD3* is significantly hypermethylated in triple negative breast cancer (TNBC) (13) compared to matched normal tissues. Second, SMARCD3 was shown to contribute to mesenchymal to epithelial transition (MET) of TNBC (14). As methylation of regulatory elements often leads to decreased gene expression, we ask whether *SMARCD3* silencing in TNBC contributes to the promotion of breast cancer, which could potentially be a valuable biomarker in predicting patient outcome and/or response to therapy. A multimodal approach has been adopted to address this question. First, we used a large, population-based data set to build prognostic models for predicting outcomes of patients with breast cancer and expressing high or low levels of *SMARCD3*. Importantly, our results show that expression of *SMARCD3* correlates with good patient outcomes, and low *SMARCD3* is associated with diminished survival rates. We investigated SMARCD3 protein expression in breast cancer samples, using tissue microarrays (TMAs) across a broad range of breast cancer subtypes, including both ER+ and ER-breast cancer samples. We identified frequent co-expression of SMARCD3 and ER, and colocalization of the two proteins in the nuclei of luminal cells. Through investigating the link between ER/SMARCD3 expression and localization, and the disease-free survival of patients, we have found that low-SMARCD3 is a marker of recurrence in ER+ patients but no correlation between ER and SMARCD3 expression.

In our effort to understand the role of SMARCD3 in cancer suppression, we show that SMARCD3 contributes to proper progression of the cell cycle, through its role as a repressor of p21. SMARCD3 depleted cells exhibit slow proliferation rates, yet higher sensitivity to genotoxic agents such as radiation.

Taken together, our data support a novel role of SMARCD3 as a breast tumor suppressor, likely through regulation of ER-mediated cell cycle checkpoints, and indicate that SMARCD3 nuclear expression has predictive value for survival in ER+ breast cancer patients.

## Results

### SMARCD3 is downregulated in breast cancer

Subunits of the SWI/SNF complex, as well as other chromatin remodeling complexes, are inactivated or mutated in many solid tumors (4, 15, 16). Consistent with this, others and our group identified *SMARCD3* as a potential tumor suppressor candidate gene in breast cancer (13). To investigate the possible link between expression of *SMARCD3* and breast cancer, we first examined the expression pattern of *SMARCD3* using real-time quantitative PCR in primary mammary carcinomas in normal (n=5) and tumor samples (n=40). The expression of *SMARCD3* varied greatly across control samples, and we detected a 3.8-fold decrease in *SMARCD3* expression in tumor samples as compared with control samples (Fig. 1B), which could be due to the limited number of normal samples accessed in this panel. To verify our initial finding in a larger cohort, we next analyzed *SMARCD3* expression in primary breast cancer and normal breast tissues using gene expression profile datasets from the TCGA breast invasive carcinoma cohort. Of 114 normal breast tissue samples and 1097 breast tumor samples, *SMARCD3* transcript level in tumors is significantly reduced compared with normal breast tissues (Fig. 1B). Normal matched samples were used as control to account for genetic diversity amongst patients. Tumor sample were found to have a decreased expression of *SMARCD3* when compared to healthy adjacent tissue (Fig. 1B; Fig.S1A & B).

To investigate the causes of SMARCD3 lowered expression in breast cancer, we interrogated methylation status of the SMARCD3 promoter. SMARCD3 promoter methylation levels were found significantly higher in tumor (T) than normal (N) in the TCGA Breast Invasive Carcinoma dataset (Fig.1C, D). Interestingly, paired analysis indicates frequent deregulation of the SMARCD3 promoter methylation status, between tumor and matched normal tissue. Finally, breast cancers with hypermethylated SMARCD3 promoters express significantly lower SMARCD3 transcripts than the matched normal samples. Taken together, these data indicate that although this decrease in expression is observed across breast cancer datasets (Fig 1; Fig. S1C) and can be explained by frequent SMARCD3 allelic copy number loss (Fig. S1B), a majority of silencing events might occur through methylation of the *SMARCD3* promoter (Fig. 1 C-F).

Members of the SMARCD family have been found mutually exclusive within the SWI/SNF complex and depletion of individual SWI/SNF subunits can destabilize the residual complex (17). Thus, to elucidate potential compensation between members of *SMARCD1/2/3* family in breast cancer, we also examined the level of *SMARCD1* and *SMARCD2* in the same samples. *SMARCD2*, but not *SMARCD1*, was significantly up-regulated in tumor samples (Fig. S1A), indicating that paralogous *SMARCD* subunits are not similarly deregulated in breast cancer. We next investigated possible correlation between *SMARCD1/2/3* expression. Using Spearman test, we found *SMARCD1* and *SMARCD2* are positively correlated in the TCGA primary tumor dataset (Fig. S1D(i)). Conversely, *SMARCD1* and *SMARCD3* are negatively correlated (Fig. S1D(ii)). Both correlations, although statistically significant, are weak and more work would be necessary to establish any biologically relevant interaction with the three SMARCD1/2/3 isoforms. No direct correlation was observed between *SMARCD2* and *SMARCD3*, indicating that even though both genes show inverse expression in tumor vs matched normal, their expression is not associated by monotonic relationship (Fig. S1D(iii), Spearman ρ = 0.005, p=0.881).

### SMARCD3, but not SMARCD1 or 2, has good prognostic value in breast cancer

Based on emerging evidence that SWI/SNF has tumor suppressor properties, we next investigated whether changes in *SMARCD3* expression are associated with specific patient survival probabilities. To explore this, we first performed survival analysis using Kaplan-Meier (KM) estimates on the TCGA Breast Invasive Carcinoma dataset. Cancer samples were categorized based on significant z-score deviation from the normal. Amongst the three genes, *SMARCD3* expression was found to be the most frequently altered (37.6%) compared to *SMARCD1* (23.6%) or *SMARCD2* (17.3%) (Fig. 1H).

KM estimates showed a significant difference in survival between patients based on their expression of *SMARCD3* (Fig. 1G). In particular, the median survival of patients with unaltered *SMARCD3* is 10.8 years (3945 days, n=566) compared to 8.9 years (3262 days, n=309) in tumors that had a low level of *SMARCD3*, suggesting that decreased *SMARCD3* expression is associated with decreased survival. Survival was not affected by *SMARCD1* or *SMARCD2* gene expression alterations (Fig. S1E). Association analysis of *SMARCD3* levels and clinicopathological variables revealed a strong association between *SMARCD3* expression and molecular subtypes and hormone receptors (Table 1). There was no significant association between *SMARCD3* expression with age, TNM Stage, T stage and N stage (Table 1). Tumors of the luminal A subtype mostly express normal levels of *SMARCD3*, while low *SMARCD3* expression is more frequent in the primary tumor of patients with ER and PR negative breast cancers and with invasive ductal carcinoma (Table 1). ER-tumors with copy number loss express significantly lower SMARCD3 when compared to normal, all tumor types, ER+ or total ER-tumors (Fig. S1B). We then broadened our analysis of *SMARCD3* prognostic value using Cox regression modelling. As a single continuous variable, *SMARCD3* expression, but not that of *SMARCD1* or *SMARCD2*, was significantly predictive of survival for patients with TCGA primary tumors (Table 1). Hazard ratio calculation correlates each unit of increase in *SMARCD3* expression with a predicted 21% decrease in fatalities (Table 1, HR=0.79 95% CI: 0.67-0.92, p=0.003).

**Table 1.** Single variable Cox regression analysis. Clinicopathological factors influencing overall survival of patients in the TCGA Breast invasive carcinoma dataset. CI: confidence Interval; HR: hazard ratio. P value derived from Wald statistical test

| Variable | SMARCD3 expression |  |  | p-value |
| --- | --- | --- | --- | --- |
|  | Low | Normal | High |  |
| **Age (years)** |  |  |  | 0.292 |
| < 45 | 55 (6.1) | 79 (8.7) | 6 (0.7) |  |
| ≥ 45 | 254 (28.0) | 487 (53.6) | 27 (3.0) |  |
| **PAM50** |  |  |  | < 0.001 |
| Lum A | 115 (12.7) | 333 (36.7) | 13 (1.4) |  |
| Lum B | 80 (8.8) | 109 (12.0) | 4 (0.4) |  |
| Her2 | 47 (5.2) | 21 (2.3) | 1 (0.1) |  |
| Basal | 64 (7.0) | 80 (8.8) | 13 (1.4) |  |
| Normal | 3 (0.3) | 23 (2.5) | 2 (0.2) |  |
| **ER status** |  |  |  | < 0.001 |
| Positive | 195 (22.5) | 455 (52.4) | 17 (2.0) |  |
| Negative | 96 (11.1) | 92 (10.6) | 13 (1.5) |  |
| **PR status** |  |  |  | < 0.001 |
| Positive | 167 (19.3) | 396 (45.8) | 15 (1.7) |  |
| Negative | 123 (14.2) | 149 (17.2) | 15 (1.7) |  |
| **HER2** |  |  |  | 0.00142 |
| Negative | 157 (25.6) | 301 (49.0) | 18 (2.9) |  |
| Positive | 67 (10.9) | 70 (11.4) | 1 (0.2) |  |
| **Pathological Stage** |  |  |  | 0.914 |
| Stage I | 48 (5.3) | 104 (11.5) | 6 (0.7) |  |
| Stage II | 184 (20.3) | 323 (35.6) | 19 (2.1) |  |
| Stage III | 71 (7.8) | 131 (14.4) | 8 (0.9) |  |
| Stage IV | 6 (0.7) | 8 (0.9) | 0 (0.0) |  |
| **Tumor stage** |  |  |  | 0.0648 |
| T1 | 74 (8.1) | 157 (17.3) | 7 (0.8) |  |
| T2 | 194 (21.4) | 319 (35.1) | 18 (2.0) |  |
| T3 | 27 (3.0) | 77 (8.5) | 6 (0.7) |  |
| T4 | 14 (1.5) | 13 (1.4) | 2 (0.2) |  |
| **Node status** |  |  |  | 0.528 |
| N0 | 144 (15.9) | 270 (29.7) | 17 (1.9) |  |
| N1 | 102 (11.2) | 194 (21.4) | 11 (1.2) |  |
| N2 | 42 (4.6) | 58 (6.4) | 3 (0.3) |  |
| N3 | 17 (1.9) | 41 (4.5) | 1 (0.1) |  |
| NX | 4 (0.4) | 3 (0.3) | 1 (0.1) |  |
| **Histological type** |  |  |  | < 0.001 |
| IDC | 261 (28.8) | 388 (42.8) | 24 (2.6) |  |
| ILC | 22 (2.4) | 125 (13.8) | 5 (0.6) |  |
| Other | 25 (2.8) | 53 (5.8) | 4 (0.4) |  |

### Immunohistochemistry (IHC) staining of patient samples reveals strong and specific nuclear localization of SMARCD3 in luminal cells

Given the possible prognostic value of *SMARCD3* identified in mRNA datasets, we decided to investigate whether mRNA levels correlate with SMARCD3 expression at the protein level using breast cancer TMAs. First, we used a commercial breast cancer TMA which contains 192 samples from exclusively female breast cancer cases distributed across all hormonal receptor status (ER, PR and HER2) as assessed by IHC (Fig. 2). The level of SMARCD3 expression was evaluated using IHC with anti-SMARCD3 antibody, after optimization of the antibody on cell blocks (SI text and Fig. S2A). Strong nuclear staining was found to correlate with luminal cells in normal acini (Fig. 2A), reminiscent of ER staining observed in normal breast tissue. These data are consistent with our previous observation that normal *SMARCD3* expression was associated with hormonal receptors and luminal A molecular subtype in the TCGA dataset. In tumor cells, nuclear intensity of SMARCD3 staining was found variable across patient samples, and scored negative (-), weak (+), moderate (++) or strong (+++) (Fig. 2B). Importantly, most samples were found reasonably homogeneous. Of the 144 cores in which SMARCD3 levels could be measured, 87 cases (60.4%) were scored negative, 24 cases (16.7%) weak, 25 cases (17.3%) moderate and 8 cases (5.6%) strong for SMARCD3 staining.

**Figure 2:**
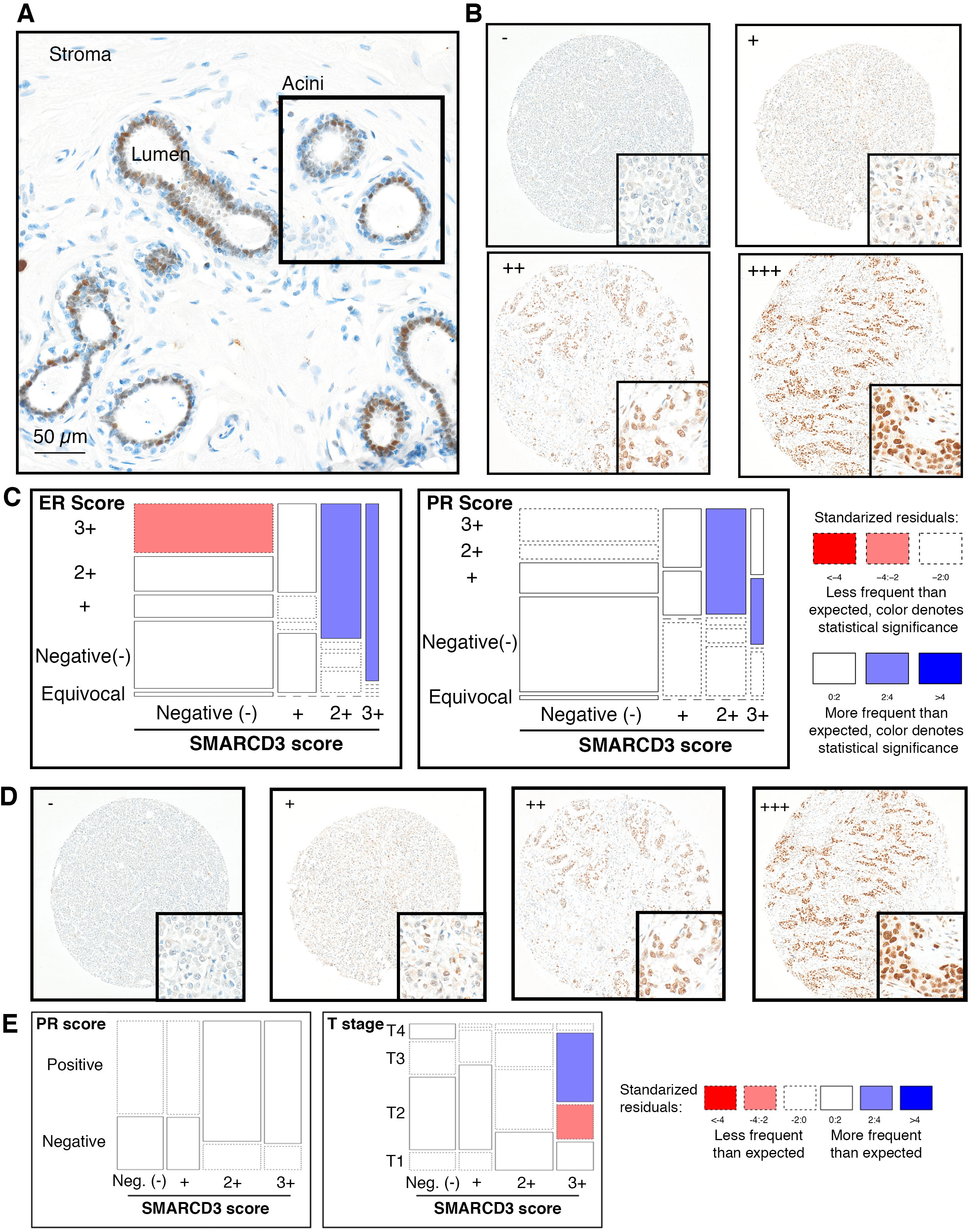
The expression of SMARCD3 in breast cancer is reminiscent of and correlate with ER IHC staining. (**A**) Expression and localization of SMARCD3 in breast cancer sample containing lobular cross-section of normal breast tissue. (**B**) Representative images of SMARCD3 immunohistochemistry staining group in tumor section of breast tissue from the BR1921b US-Biomax cohort. (-) denote absence of staining, (+) weak nuclear staining, (++) moderate nuclear staining and (+++) strong nuclear staining. **(C)** Mosaic plot of variables (ER and PR status inferred by IHC) significantly associated with SMARCD3 expression in the US-Biomax breast cancer cohort. The sizes of tiles correspond to the frequency of breast cancer patients that fall within each category. The color and shading represent the significance and strength of the deviation from the null hypothesis that the hormone receptor status and SMARCD3 intensity are randomly distributed: blue indicates that a category is overrepresented; red is underrepresented.

Contingency tables were built to investigate the relationships between SMARCD3 nuclear scores and clinicopathological variables and mosaic plots were used to investigate the direction of significant deviation (Fig. 2B). SMARCD3 protein levels were significantly associated with the IHC status of ER and PR (Table S1, p<0.001). A direct correlation was observed between the level of SMARCD3 and ER (Fig. 2B), and between moderate/strong SMARCD3 level and strong PR signal. Interestingly, the correlation between SMARCD3 expression and ER level of expression is not complete, as negative SMARCD3 tumors expressed the full range of ER expression (Table S1, Fig. 2B).

As IHC validated our previous mRNA analyses, we wondered whether SMARCD3 protein level could be used to predict patient outcome. To answer this query, we made use of a cohort recently published (18), which contains 229 ER+, HER2-breast cancer patients with axillary nodal metastatic disease, with median followup time of 4.9 years. Relapse was used as a surrogate for breast cancer survival and occurred in 46 patients (20.1%; see (18)) with a median relapse-free time of 7.7 years (SD 3.8 years). SMARCD3 expression using IHC could be assessed in 207 tumor cores and SMARCD3 nuclear staining intensity (Fig. 2C(i)) was recorded as negative in 55 cases (26.6%), weak in 39 cases (18.8%), moderate in 69 cores (33.3%) and strong in 44 samples (21.3%). Chi-square analysis indicated a significant direct relationship between SMARCD3 level and PR expression (Table S2, p=0.018). Mosaic plots did not indicate a significant correlation between specific categories, however negative and weak SMARCD3 associates with negative PR expression, whereas moderate and strong SMARCD3 expression tend to associate with positive PR expression (Fig. 2D). Significant differences in breast cancer relapse were observed with patients grouped by low (-/+) SMARCD3 (n=94) *vs* high SMARCD3 (++/+++; n=113), (Fig. 2D(ii) and (iii)). KM plots (Fig. 2D) and hazard ratio calculation (Table S3) indicated that high-SMARCD3 expression (as defined by IHC: ++/+++) halves the risks of relapse compared to low-SMARCD3 (as defined by IHC: -or +) patients (HR=2.11 95% CI: 1.10-4.06, p=0.024).

### Cells depleted for SMARCD3 are deficient for DNA damage repair

The DNA damage response, comprising an intricate network of DNA damage checkpoints and damage repair mechanisms, allow cells to maintain their genomic integrity. For this reason, many DNA repair proteins are suppressors of breast tumors (19). Given the potential protective role of SMARCD3 in ER+ breast cancer, we wondered whether this tumor suppression activity could be attributed to a DNA damage repair role. We monitored γH2AX immunofluorescence as a surrogate for DNA double-strand breaks (DSBs), and used γ-ray irradiation as the DSB inducing agent, we observed an accumulation of γH2AX foci in cells depleted for SMARCD3 when compared to control cells. Interestingly, γH2AX foci form in SMARCD3 depleted cells even in the absence of genotoxic stress (t=0, Figure 3A (i) and (ii)). In addition, following irradiation, the resolution of γH2AX foci is delayed in the absence of SMARCD3 when compared to control cells (Fig. 3A,B). To confirm that SMARCD3 depleted cells are impaired for DNA damage repair pathways, we examined survival using colony formation upon irradiation (Fig. 3C). SMARCD3 depletion led to radiation hypersensitivity, noticeable even at low dose (2Gy), indicating that cells with low SMARCD3 expression are indeed impaired for DNA damage repair. EdU incorporation in cells expressing SMARCD3 or depleted by shRNA, and treated with 6 Gy irradiation, showed a minor reduction of EdU staining upon SMARCD3 depletion compared to the control (Fig. 3C(iii)). Interestingly, investigation of *SMARCD3* expression level the Homology Directed Repair (HDR) signature (e.g. expression of other genes whose expression decreases in response to DNA damages) indicated a direct correlation between *SMARCD3* and the (HDR) signature (Fig. 3D) (20). For each cohort (low-expressing and high-expressing SMARCD3), we investigated the following markers: Number of telomeric <u>a</u>llelic imbalances (NtAI); <u>L</u>arge-<u>s</u>cale <u>s</u>tate transitions (LST); Number of genomic segments with Loss Of Heterozygozity (HRD-LOH) and finally the total HRD score, which is the sum of each individual marker.

**Figure 3.**
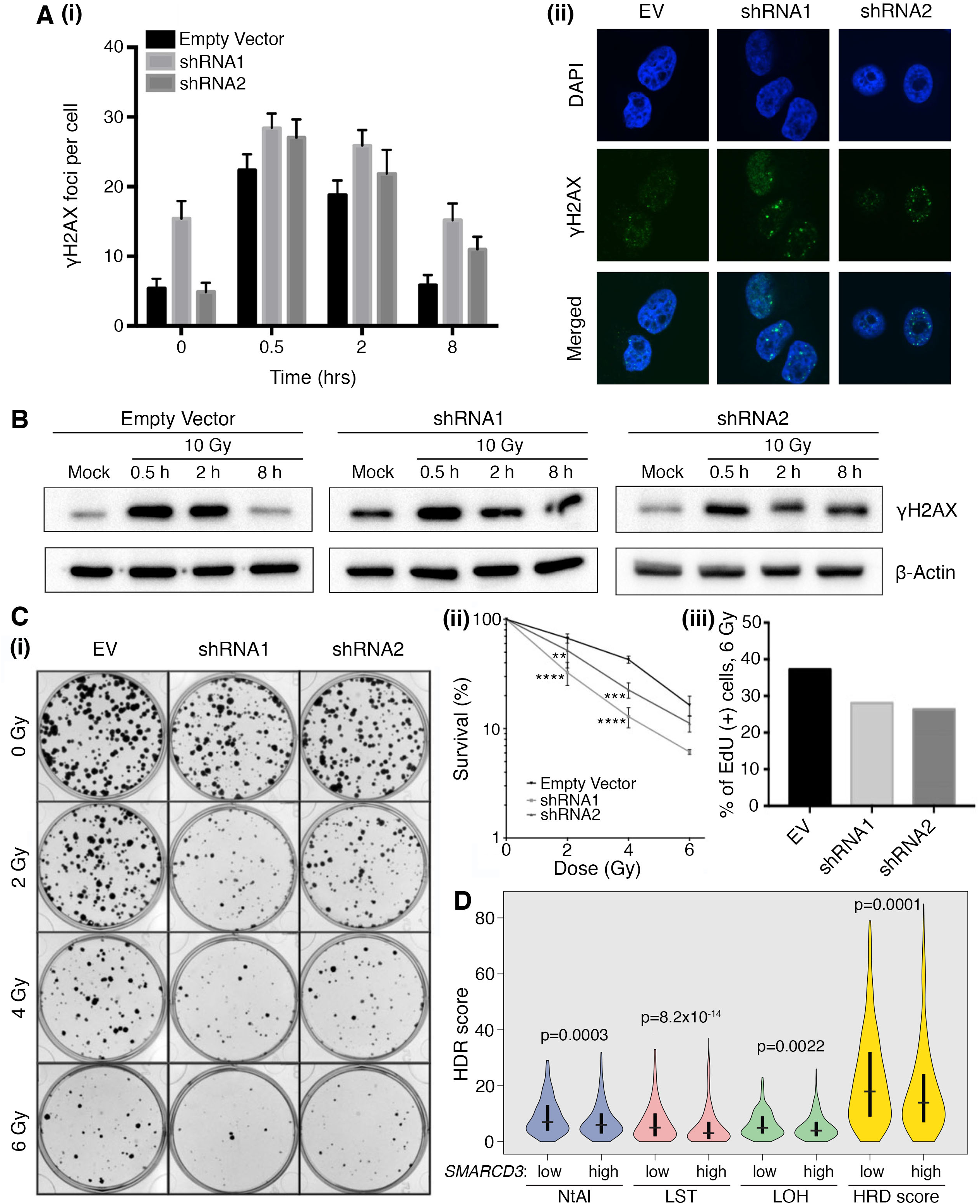
The expression of SMARCD3 in breast cancer cell lines. **(A)** (i) γH2AX foci were quantified at 0, 0.5, 2 and 8 h post 2 Gy irradiation (ii) Representative images from immunofluorescence assay of Empty Vector HeLa cells and SMARCD3 depleted HeLa cells at 8 h post 2 Gy irradiation treatment and stained with DAPI and probed for γH2AX protein. **(B)** Western analysis γH2AX in of Empty Vector HeLa cells lysate and SMARCD3-knockdown HeLa cells lysate at 8 hours post 10 Gy irradiation treatment using an anti-γH2AX antibody and anti-β-actin as loading control. **(C) (i)** Representative colony forming ability of HeLa *SMARCD3* knockdown and empty vector cells in response to gamma-radiation. (ii) The percentage survival of Empty Vector HeLa and SMARCD3-knockdown HeLa cell lines in response to an increasing dose of gammaradiation in Grays (Gy). Data are shown as mean +/ SD of three independent experiments. (iii) The Percentage of EdU positive HeLa *SMARCD3* knockdown and empty vector cells 24 h post-treatment with 2 Gy gamma-irradiation. Results were analyzed by one-way ANOVA with Dunnett’s multiple comparisons test correction. ** adjusted P-value < 0.01; *** adjusted P-value< 0.001. **(D)** HDR score analysis of TCGA ER+ breast cancer patients. Median SMARCD3 expression level was split at the median into SMARCD3-low and SMARCD3-high cohorts. The following markers of homologous recombination deficiency (HRD) are plotted for each cohort: NtAI: Number of telomeric <u>a</u>llelic imbalances; LST: <u>L</u>arge-<u>s</u>cale <u>s</u>tate transitions; HRD-LOH: Number of genomic segments with LOH; HRD score: The sum of each aforementioned score. Statistics: Mann-Whitney *U* test.

This analysis shows that the number of each type of homologous recombination deficiency (HRD) and the total HRD score is significantly higher in *SMARCD3*-low than in *SMARCD3*-high ER+ samples.

### Depletion of SMARCD3 perturbs cell cycle progression and overall proliferation

Uncontrolled proliferation is a key characteristic of cancer cells and catalytic subunits of the SWI/SNF complex are known to impact upon proliferation and senescence (21, 22). Several DNA repair pathways are subject to cell cycle control. For instance, resection of DSB ends, which is a critical first step in homologous recombination, is normally repressed in the G1 phase of the cell cycle. We investigated whether the DNA damage repair deficiency in and reduced survival of SMARCD3 depleted cells might have stemmed from perturbations in cell cycle progression. Stable knockdowns were generated in HeLa, as well as in triple negative breast cancer cells MDA-MB-231 and MDA-MB-468 (Fig. 4 and S3), which all have high baseline expression of SMARCD3, using two independent short-hairpin RNAs or an empty lentiviral vector (EV) as control (Fig. 4 and S3).

**Figure 4.**
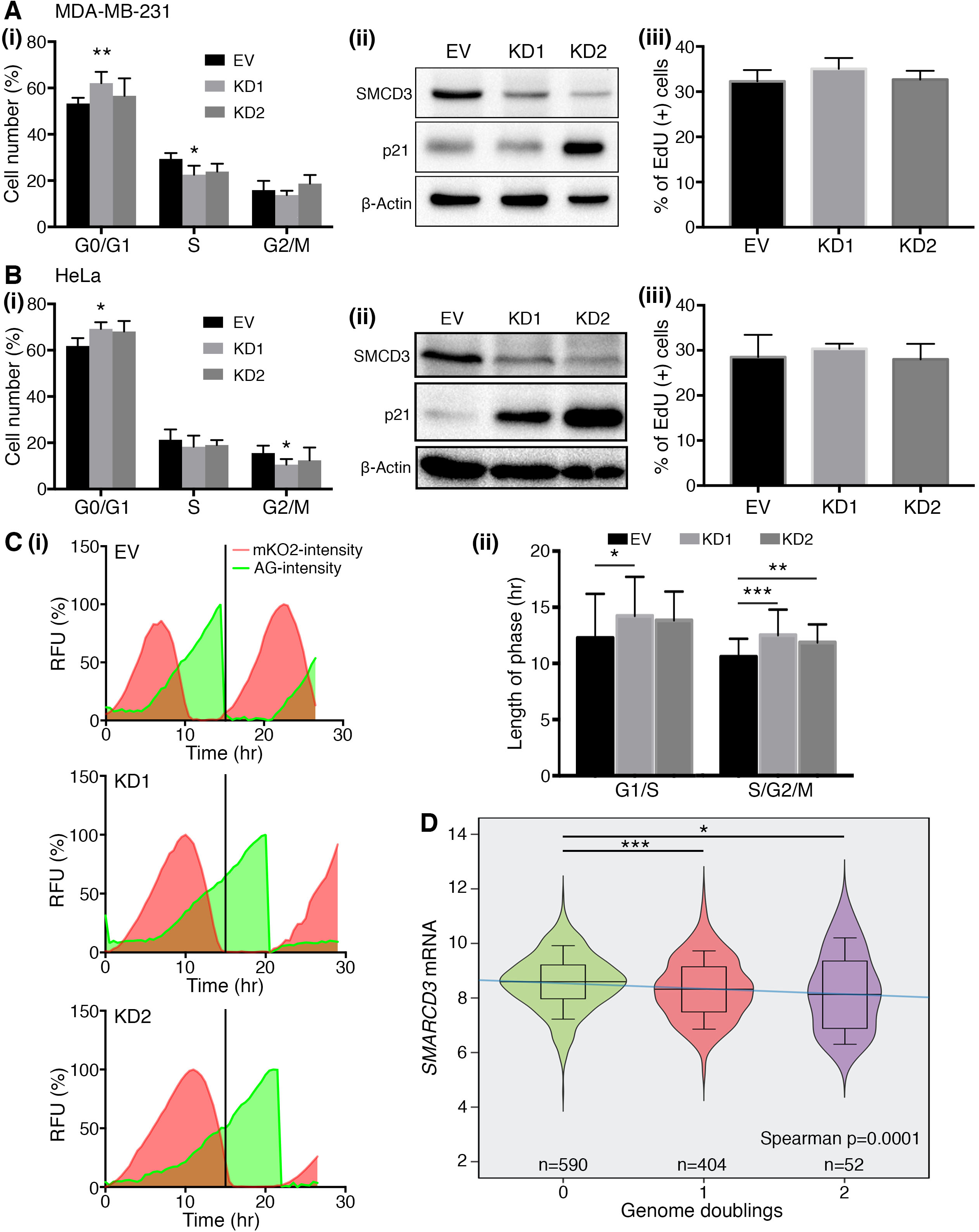
Knockdown of *SMARCD3* promote genome instability. (**A**), **(B)** The following were investigated in MDA-MB-231 (A) and HeLa (B): (i) Cell cycle distribution by PI staining. The corresponding percentage of cells remaining in G0/G1, S or G2/M is plotted as mean +/-SD for three independent technical replicates. Significance assessed using t-test. (ii) Western blot of SMARCD3 and p21 proteins in SMARCD3 depleted cells and controls (iii) Percentage of EdU positive cells depleted for *SMARCD3* or controls following 1h EdU incorporation. (**C**) Representative single-cell time series tracks of mKO2-hCdt1 and mAG-hGeminin arbitrary relative fluorescence units (RFU) percentages over time in HeLa-FUCCI cells transduced with EV control, or *SMARCD3* shRNAs. (ii) Average duration of individual cell cycle phases is plotted (mean ± SD for 30 individual cell tracks). Significance determined using unpaired t-test. **(F)** Violin plot comparing *SMARCD3* gene expression distribution across breast cancer samples that have undergone none (green), single (red) or double whole-genome doubling (purple). Significance determined by Mann Whitney *U* tests, blue line denotes significant inverse correlation between *SMARCD3* expression and genome doubling (Spearman’s rank correlation P value). In all panels * p<0.05, ** p<0.01, *** p<0.001, **** p<0.0001, ns = non-significant.

First, we followed MDA-MB-231 and MDA-MB-468 growth rates by live-cell imaging for over 96 hours. MDA-MB-231 (Fig. S3A) and MDA-MB-468 (Fig. S3B) cells depleted for SMARCD3 exhibited lower proliferation rates than control cells. We used PI staining of the DNA and flow cytometry to investigate the cell type profile of HeLa cells and the triple negative breast cancer cell line (MDA-MB-231) depleted of SMARCD3. With shRNA1, cells depleted for SMARCD3 showed a modest accumulation in G1 when compared to controls, and a slight decrease in the numbers of cells entering S phase (Fig. 4A, 4B(i); raw data shown in Fig. S3C). To investigate G1/S transition, we measured the level of cyclin-dependent kinase inhibitor p21 by Western blotting. This revealed an inverse correlation between SMARCD3 and p21, with strong accumulation of p21 in response to SMARCD3 depletion (Fig. 4A, 4B(ii)). Importantly, the level of p21 does not correlate with the cell proliferation rate, indicating that cells depleted for SMARCD3 might not respond to activated checkpoints. Consistently, incorporation of EdU indicated no noticeable difference between cells depleted for SMARCD3 or the control (Fig. 4A, 4B(iii)), which hints at normal DNA synthesis. Subtle variations in the cell cycle can be overlooked when using populations of cells. We used the FUCCI system (23) and live-cell imaging to investigate the behavior of single cells. While the ratio of cells in each phase of the cell cycle does not change, individual phases of the cell cycle are longer in cells depleted for SMARCD3 than in control cells (Fig. 4C and movies M1, M2). In addition to a longer S-phase, the S/G2 transition is compromised in cells depleted for SMARCD3, and we observed a higher incidence of cells undergoing endoreplication, where cells initiate a new cycle of DNA synthesis instead of entering mitosis. This results in whole-genome doubling, a phenomenon estimated to occur in 45% of breast cancers (24). Consistent with our *in vitro* data, we find that SMARCD3 expression is significantly lower in breast tumors that had undergone one or two whole-genome doublings (Fig. 4D).

## Discussion

Here, we provide evidence that *SMARCD3* is downregulated in malignant breast tissue, which may indicate a tumor suppressor function of *SMARCD3* in breast cancer. We used three cohorts of breast cancer patients with various clinicopathological features to reach this conclusion. In all three, we observed significant downregulation of *SMARCD3* in breast cancer relative to normal breast tissue. This indicates that besides mutations and copy number loss targeting the *SMARCD3* gene locus, transcriptional inactivation of *SMARCD3* gene, such as by promoter methylation, might contribute to the cancer phenotype. SMARCD1, 2 and 3 are three isoforms that can each be incorporated into the BAF chromatin remodeling complex. However, we have demonstrated that the expression of the three isoforms is not correlated either positively or negatively, indicating that they probably do not compensate for each other in cells when one is absent or downregulated. Interestingly, *SMARCD3* was the only subunit that exhibited a strong link with breast cancer with prognosis value, with low *SMARCD3* correlating with worse patient outcome. On average, the survival of patients with unaltered *SMARCD3* is 673 additional days (1.9 years) when compared to patients with low *SMARCD3*, indicating a dramatic difference in outcome between these two patient groups. Thorough analysis of confounding factor is warranted to validate the predictive value of *SMARCD3*. In that regard, our histological investigation of SMARCD3 indicated a possible link between SMARCD3 and hormone receptors. In breast cancer, estrogen can be a major driver of cell proliferation, as well as an efficient means for therapeutic intervention. Interestingly, we found that in normal breast tissue, SMARCD3 localizes in the nuclei of luminal cells, which is the same localization pattern as the estrogen receptor (ER). Similarly, the staining of SMARCD3 in ER+ breast cancer is reminiscent of that of ER, although expression of SMARCD3 and ER do not fully correlate. While high SMARCD3 is only found in strong ER+ samples, where both proteins could contribute to increased proliferation of the tumor, low SMARCD3 was observed across the full range of tumors, irrespective of ER expression level.

Conversely, both high- and low-SMARCD3 expression correlated with the expression of Progesterone Receptor (PR). Although more studies are required to fully understand the link between SMARCD3 and both ER and PR hormone receptors, the inclusion of SMARCD3 as a prognostic marker, together with PR and HER2 status, may help stratify ER positive breast cancer patients and aid in treatment decision making.

While SMARCD3 depleted cells appear to exhibit slower division rates as measured by live-cell imaging and accumulate high levels of G1/S checkpoint proteins, further investigation by live-cell imaging and the FUCCI system demonstrated that cells depleted for SMARCD3 might undergo DNA synthesis and cell cycle progression like the WT counterpart, only failing in the G2/M transition and undergoing endoreplication as a result. This finding is consistent with our observation that breast tumors that have undergone whole-genome doubling express significantly lower levels of *SMARCD3* than tumors that have not. It also suggests that high risks of genomic instability do not always correlate with increased mitotic index. While we expected that SMARCD3 depletion would impair the normal progression of the cell cycle by limiting DNA replication through its role in chromatin remodeling, we found that replication is proficient. However, termination of replication and then mitosis are compromised. Our findings that SMARCD3 levels directly impact cell cycle progression by regulating p21 provides an additional clue to explain the role of SMARCD3 in carcinogenesis. Although the absence of SMARCD3 correlates with a low mitotic index, tumors expressing low levels of SMARCD3 could be considered aggressive given a heightened risk of recurrence and lack of response of these cells to p21. Our work provides evidence that low-SMARCD3 expression in ER+ tumors predicts poor response to the current standard of care, and patients with such tumors might benefit more from aggressive or personalized treatments. As a subpopulation of SMARCD3 cells accumulates in G0, it is easy to speculate that cells expressing low SMARCD3 levels might stay dormant and delay their re-entry into the cell cycle, accounting for some of the recurrence observed in poor-prognosis ER+ patients. In the future, it will be interesting to test whether SMARCD3 inactivation could provide a means to sensitize ER+ cells to genotoxic chemotherapy or radiotherapies.

## Acknowledgments

ED and PHGD are recipient of National Breast Cancer Fellowships. RT and BDLP are recipient of the Princess Alexandra Research Foundation. This work was funded by ECR13-04 (NBCF) and Cancer Council Queensland (CCQ). The Mays cancer center is supported by a NCI Cancer Center Support Core Grant P30 CA054174.

## Methods

### Cell lines culture

MDA-MB-231, MDA-MB-468 and HeLa were obtained from ATCC and cultured in DMEM supplemented with 10% fetal bovine serum (FBS).

### Lentiviral particles production and cell infection

Replication incompetent lentivirus were produced in HEK 293T cells co-transfected by mixing 5 µg of either pLKO.1 Empty Vector or pLKO.1 shRNA targeting *SMARCD3* with 6 µg Lenti-vpak packaging kit components (OriGene) and 33 µL of transfection reagent (OriGene) in 1.5 mL OptiMEM, incubated for 20 min at RT. This transfection mixture was added onto 10 mL of HEK 293T cells at 2×10^5^ cell/mL in fresh culture medium was transferred. Culture medium was replaced 24h post-transfection and viral supernatants were harvested twice at 48 and 72h and combined. Viral supernatants were clarified by centrifugation at 500 *g*, filtered through a 0.45 μm PES and stored at −80°C. Lentiviral transduction was performed with 5×10^5^ cells seeded into a T25 flask in media supplemented with 4 μg/mL polybrene. Viral supernatant (500 µL) was added onto the cells and they were then incubated for 24h. 72h post-transduction, cells were selected with Puromycin (for pLKO) at 1 µg/mL. After two weeks of Puromycin treatment cells were considered selected. Knockdown efficiency of *SMARCD3* was determined by Western Blotting and/or qRT-PCR as described below. Cells transduced with FUCCI plasmids were sorted by FACS to select homogenous positive cells populations.

### RNA extraction and cDNA preparation

Total RNA was extracted from cells in exponential growth on 10 cm dishes. Cells were washed twice with PBS and 2 mL of TRIzol (Invitrogen™) was added on each plate. 500 µL of cell extracts were transferred into microfuge, chloroform extracted and nucleic acid was precipitated with isopropanol/ethanol. RNA was dissolved in RNase-free water and stored at −80°C. RNA yield and purity were evaluated on a NanoDrop and agarose gel electrophoresis. The reaction mixture for reverse transcription was prepared as previously described (26). Briefly, 800 ng total RNA resuspended in RNase-free water was combined with 2.5 μM random hexamer and 500 μM RNase-free dNTPs (both Invitrogen™), heated at 65°C for 5 min, cooled on ice for at least one minute while adding 4 µl SuperScript™ III reaction buffer, 5 mM dithiothreitol, 40 U RNase out RNase inhibitor and 200 U SuperScript™ II RT (all from Invitrogen™) to each sample, except “no RT” control tubes. The mixture was incubated at 25°C for 5 min, at 50°C for 60 min and finally at 70°C for 15 min in a thermal cycler (Eppendorf). cDNA used to generate Figure 1B were sourced from OriGene (BRCT102).

### Real-time quantitative PCR (RT-qPCR)

RT-qPCR was performed using Taqman^®^ technology in 10 μL reaction volume. The reaction mixture was prepared as follow: 2 μL of cDNA, 5 μl Taqman^®^ Universal Master Mix (Invitrogen™), 2.5 µL of water and 0.5 μL of Taqman^®^ probe: human *SMARCD3* (Invitrogen™ Cat. # 4331182) and human *18s* (Invitrogen™ Cat. # 4331182) was used as the reference endogenous control. Taqman^®^ probes were carefully picked to span exon-exon junction and thus, specifically target cDNA. The RT-qPCR assays were completed using the ABI ViiA™ 7 sequence detection system (Applied Biosystems^®^) with the following parameters: initial denaturation 50°C for 2 min then 95°C for 10 min followed by 40 cycles of 95°C for 15 seconds and 60°C extension for 1 min. Fluorescence was quantified at the end of each cycle. The real-time PCR data was analyzed using the ΔΔCt method.

### Western blot analysis

Cells were pelleted and lysed using RIPA buffer (10 mM Tris-Cl (pH 8.0), 1 mM EDTA, 1% Triton X-100, 0.1% sodium deoxycholate, 0.1% SDS, 140 mM NaCl, 1 mM PMSF). Lysates were sonicated and protein concentration was determined using BCA protein assay (Sigma). 30 μg of total proteins were loaded on 12% SDS–PAGE gel and separated by electrophoresis. Protein were transferred on PVDF membrane, dried and blotted overnight in 5% BSA TBS-T at 4 °C using primary antibodies (Abcam anti-SMARCD3: ab171075; Santa Cruz anti-β Actin (C-4): sc-47778; Millipore anti-γH2AX (Ser139): 05-636; Cell Signaling anti-p21WAF^1^/CIP^1^ (12D1): 2947). Afterwards, membranes were incubated with secondary antibodies for one hour at room temperature and developed using Clarity ECL Detection Kit (Bio-Rad).

### Bioinformatics

All available TCGA RNA-seq V2 data and clinical annotations were downloaded from the UCSC Xena browser. Data from the METABRIC dataset was obtained from cBioPortal and Oncomine. Both TCGA and METABRIC research data are publicly available and all patient information is de-identified. For Kaplan-Meier analysis, gene expression was converted into discrete variable to categorize patient based on *SMARCD3* expression. *SMARCD3* expression mean and standard deviation from the normal sample distribution were used to score sample as a z-score. Each sample with a *SMARCD3* expression included within 2 standard deviation from the mean was score as unaltered, while samples with 2 standard deviation away from the mean were score as either low or high (i.e. p < 0.05). The number of whole-genome doublings in each TCGA breast cancer sample was determined using ABSOLUTE (27).

### Statistical analysis

Statistical analysis of gene expression in cancer sample was carried out in GraphPad Prism. Survival analysis was carried out in R version 3.4.2 (R Development Core Team; www.r-project.org) Kaplan-Meier estimate, log-rank test and curves were built with Survival and surviminer package (v. 2.41-3 and v. 0.4.1.99, respectively). Cox proportional hazard models were built with the rms-package (v. 4.3-1). Single Cox’s proportional-hazards regression were fitted on gene expression and clinical variables. The hazard ratio and its 95% CI were derived from these models and significance of the results were determined by Wald Statistical test. For the HDR score analysis, TCGA ER+ breast cancer patients were split by median SMARCD3 expression level, referred to as SMARCD3_low and SMARCD3_high cohorts. Then the following measures for homologous recombination deficiency (HRD) are compared in the cohorts (i) NtAI: The <u>n</u>umber of telomeric <u>a</u>llelic imbalances, (as per (28)); (ii) LST: <u>L</u>arge-<u>s</u>cale <u>s</u>tate transitions, (as per (29)); (iii)HRD-LOH: Number of genomic segments with LOH, (as per (30)) (iv) HRD score: The sum of each of the above, (as also per (31)). Statistics: Mann-Whitney *U* test.

### Immunohistochemistry (IHC)

Antigen retrieval was performed with the Ventana Discovery ULTRA Staining module, using Discovery CC1 for 32 min, on TMA from US Biomax (1921b) or Mater patient cohort (18). Primary immunostaining was performed using antibodies against SMARCD3 (Abcam ab1711075) and Ventana antibody dilution buffer for 36 min at 36°C. Secondary immunostaining used an anti-rabbit horseradish peroxidase conjugated antibody and immune complexes were visualized using diaminobenzidine tetrahydrochloride, followed by counter-stain with hematoxylin II for 8 min. The scoring of the immunostained TMAs was performed by two independent investigators based on the SMARCD3 nuclear intensity level, under the guidance of a pathologist.

### Cell proliferation assays

Automated live cell imaging was conducted using an IncuCyte Zoom (Essen BioScience, Ann Arbor, MA, USA) live-cell imaging system. Cells were seeded onto a 96-well plate at 3×103 cells per well. Phase contrast images of cells were acquired every two hours and proliferation was measured as a percentage of confluency obtained from the IncuCyte cell proliferation analysis module. The proliferation data was analysed in GraphPad Prism v7.0. A non-linear exponential curve was fitted and the population doubling time and R2 goodness-of-fit constant were extracted from the non-linear exponential growth curve analysis module in GraphPad Prism v7.0.

### Clonogenic assay

Cell death in response to irradiations was assayed by measuring colony formation with and without increasing doses of gamma irradiation (0, 2, 4 and 6 Gy) treatment using a Gammacell® 40 Exactor (Best Theratronics Ltd, Kanata, ON, CA). 300 cells per well were seeded on a 6-well plate and incubated for 8 h before being treated with increasing dose of gamma radiation (2, 4 and 6 Gy) using a Gammacell® 40 Exactor (Best Theratronics Ltd, Kanata, ON, CA). Culture medium was replaced every three days, and cells were incubated for 10-11 days or until surviving individual cells at T0 had divided sufficiently to form a visible colony (estimated to be 60 cells). Cells were fixed with methanol-acetic acid (3:1) and stained with 0.4% crystal violet. Excess dye was removed by immersing the plate in clean water twice. The 6-well plates were imaged using a chemidoc (Bio-Rad, Hercules, CA, USA) and the number of colonies was counted in CellProfiler using an automated analysis pipeline. Relative colony formation (%) as measured by the number of colonies from treated well divided by the number of colonies in the untreated control and plotted.

### Indirect immunofluorescence

Stable cell lines expressing shRNA or control plasmid were grown on coverslips for 24h and treated with 4 Gy γ-rays. Cell nuclei were pre-extracted with nuclear extraction buffer (10 mM PIPES (pH 6.8), 100 mM NaCl, 300 mM sucrose, 3 mM MgCl_2_, 1 mM EGTA (pH 8.0), 0.5% Triton X-100) for 2 min at RT and fixed with 4% paraformaldehyde (PFA) for 10 min at 4°C. Nuclei were blocked in 5% BSA/0.3% Triton X-100 for 2h at RT, immunoblotted with a primary antibody (Millipore anti-γH2AX (Ser139): 05-636) in 1% BSA/0.3% Triton X-100 for 2h at RT, followed by secondary antibody for 2h at RT. DNA was stained with ProLong Gold/Diamond Antifade Mountant with DAPI. Number of cells with nuclear foci were quantified using CellProfiler.

### Cell cycle progression (flow cytometry)

Cells (asynchronized population) were collected and fixed with 70% ethanol at −20°C for at least 24h. DNA was stained with 100 µg/mL RNase A (QIAGEN) and 150 µg/mL propidium iodide (PI) for one hour at RT. Data were collected using a CytoFLEX Flow Cytometer (Beckman Coulter) and results were analyzed with FlowJo. 10,000 events were collected and aggregated cells were gated out.

### Click-iT™ EdU

Control and knockdown cells (4.0 x 10^4^ cells/well) were seeded in 12-well plates with glass coverslips for 24h. To label the cells (according to manufacturer’s protocol), 5-ethynyl-2’-deoxyuridine (EdU) (Life Technologies) was added to a final concentration of 10 µM and incubated for 90 min at 37°C. Cells were fixed with 4% (PFA) for 10 min at RT, followed by permeabilization with 0.3% Triton X-100 for 20 min at RT. Click-iT™ Reaction Cocktail with Alexa Fluor^®^ 488 (Life Technologies) was added to fixed cells and incubated for one hour at RT. Nuclei were stained with ProLong® Diamond Antifade Mountant with DAPI (Invitrogen™). EdU-stained cells were quantified using CellProfiler.

**Sup. Table 1 SMARCD3 protein expression classified by clinicopathological variables** (hormone receptor status; stage and grade) in 144 patients where staining was observed from the BR1921b US-Biomax cohort.

**Sup. Table 2 Relative representation of SMARCD3 IHC scores across clinicopathological variables** in 206 primary tumors of patients from the ER+, HER2-Breast Cancer Research group cohort.

**Sup. Table 3 Single variable Cox regression analysis**. Clinicopathological factors influencing overall survival of patients in the ER+, HER2-Breast Cancer Research group cohort.

**Sup Fig 1 (A)** RNAseq analysis of *SMARCD1/2/3* gene expression in TCGA breast invasive carcinoma dataset in matched tumour and normal samples (paired t-test; **** P <0.0001, ns = non-significant). (**B**) Copy number variation (CNV). (**C**) **(D)** Scatter plot of primary tumor *SMARCD1/2/3* gene expressions with strength and direction of the correlation indicated by parametric (Pearson) and non-parametric (Spearman) coefficient and P values for significance obtain from GraphPad Prism. (i) SMARCD2 expression versus SMARCD1 expression (ii) SMARCD3 expression versus SMARCD1 expression (iii) SMARCD3 expression versus SMARCD2 expression. (P * < 0.05, ** < 0.01) (**E**) Kaplan Meier curves of overall survival of patient stratified using gene high, normal or low expression (i) SMARCD1 expression (iii) SMARCD2expression in the primary tumor. P values derived from the log-rank test.

**Sup Fig 2 (A)** (i) optimization of anti-SMARCD3 Immunocytochemistry staining using HeLa WT cells embedded in a paraffin block (ii) immunocytochemistry of HeLa cells control and SMARCD3-depleted cells embedded in a paraffin block. **(B)** (i) Representative images of SMARCD3 immunohistochemistry staining group in tumor section of breast tissue from the Breast Cancer Research group cohort. (ii) Kaplan Meier curve of relapse-free survival of patient classified using SMARCD3 IHC scoring by pooling negative and weak scored tumors into a Low SMARCD3 group and moderate and strong scored tumors into a High SMARCD3. P value were derived from log-rank test.

**Sup Fig. 3** shRNA-mediated knockdown of SMARCD3 in MDA-MB-231 (**A**) and MDA-MB-468 (**B**). (i) Knockdown efficiency was analyzed by Western blot. (ii) Control and SMARCD3 knockdown cells were incubated at 37°C 5% CO2 in the IncuCyte live-cell imaging system as described in the methods section. The time course of cell proliferation was measured by live-cell imaging (pictures at 2-hour time intervals) as a percentage of confluence. **(C)** Cell cycle distribution of SMARCD3-depleted cells and Empty-vector control in (i) MDA-MB-231 and (ii) HeLa cells.

