## Supplementary material for "The chromatin remodeler SMARCD3 regulates cell cycle progression and its expression predicts survival outcome in ER+ breast cancer": 3 Sup Figures and 3 sup tables

### Supp. Figure 1_Tropee_2019

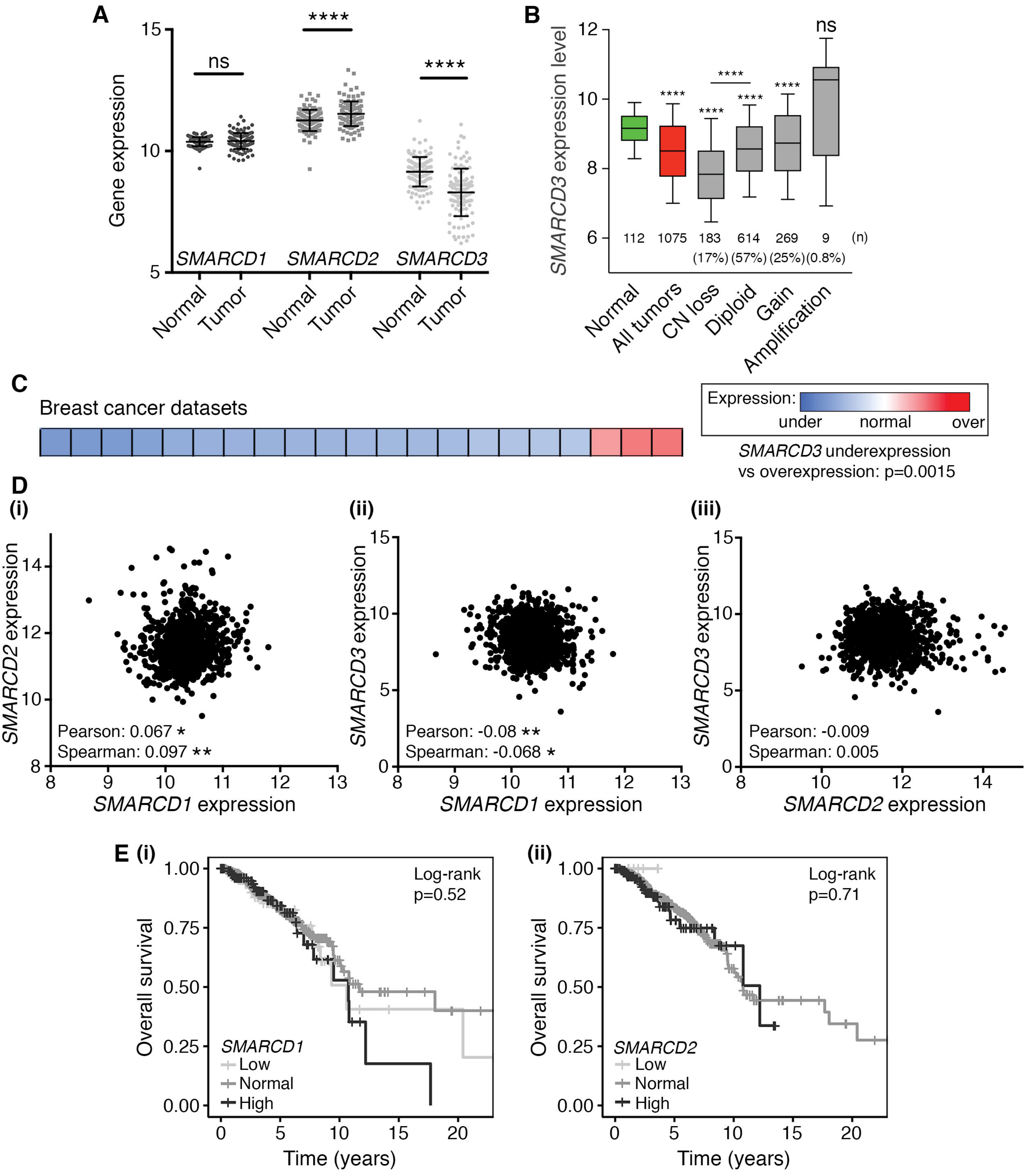

### Supp. Figure 2_Tropee_2019

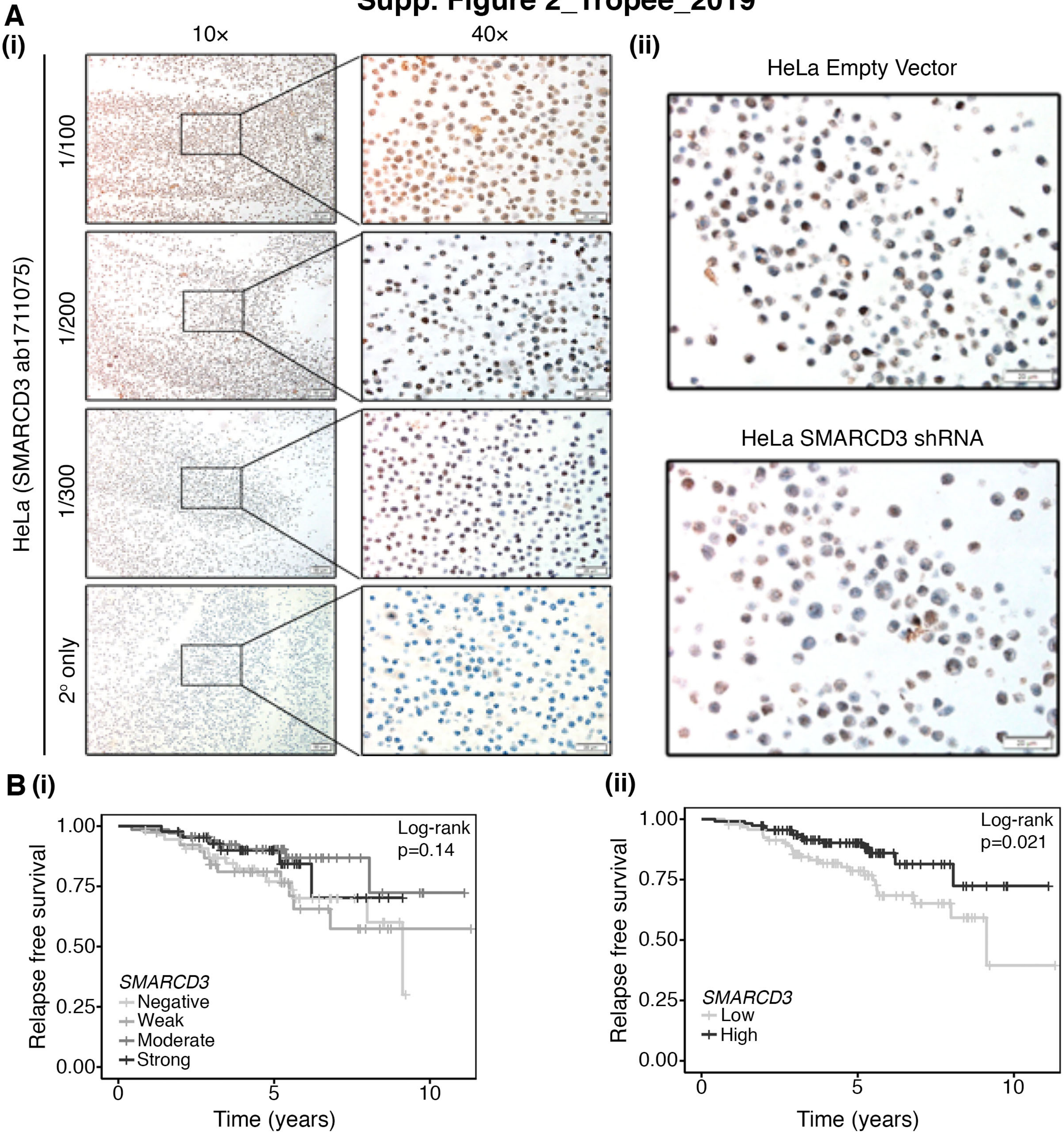

### Supp. Figure 3_Tropee_2019

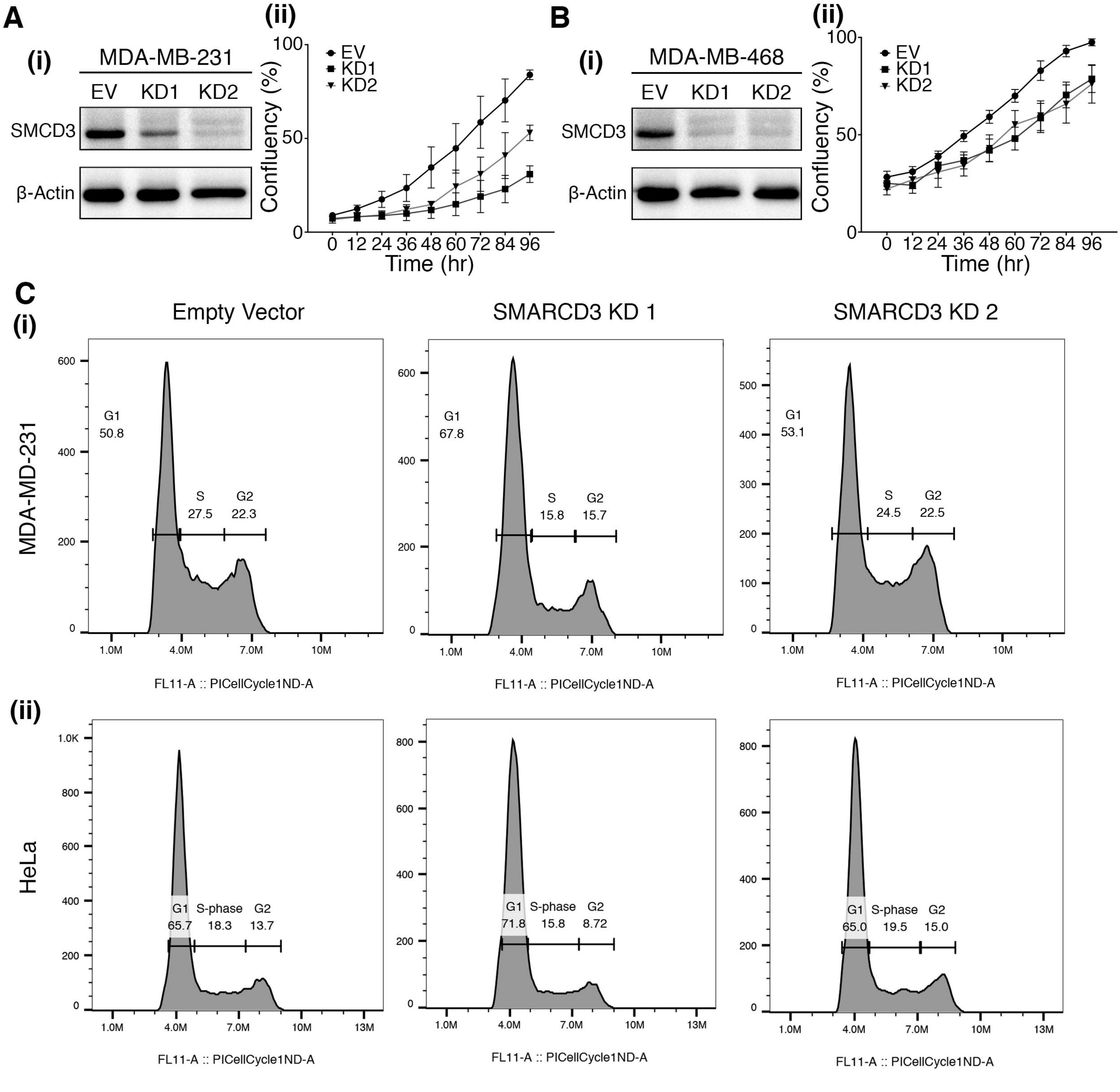

#### Supplementary Table 1_Tropee_2019

| | SMARCD3 score | | | | |
|---|---|---|---|---|---|
| | **-** | **+** | **++** | **+++** | **p-value** |
| **Age (years)** | | | | | 0.421 |
| < 45 | 23 (16.0) | 5 (3.5) | 10 (6.9) | 3 (2.1) | |
| ≥ 45 | 64 (44.4) | 19 (13.2) | 15 (10.4) | 5 (3.5) | |
| **ER** | | | | | **0.001** |
| Equivocal | 2 (1.4) | 0 (0.0) | 0 (0.0) | 0 (0.0) | |
| - | 33 (22.9) | 8 (5.6) | 3 (2.1) | 0 (0.0) | |
| + | 11 (7.6) | 1 (0.7) | 2 (1.4) | 0 (0.0) | |
| ++ | 17 (11.8) | 3 (2.1) | 1 (0.7) | 0 (0.0) | |
| +++ | 24 (16.7) | 12 (8.3) | 19 (13.2) | 8 (5.6) | |
| **PR** | | | | | **< 0.001** |
| Equivocal | 2 (1.4) | 0 (0.0) | 0 (0.0) | 0 (0.0) | |
| - | 47 (32.6) | 10 (6.9) | 7 (4.9) | 2 (1.4) | |
| + | 15 (10.4) | 0 (0.0) | 2 (1.4) | 0 (0.0) | |
| ++ | 7 (4.9) | 6 (4.2) | 1 (0.7) | 3 (2.1) | |
| +++ | 16 (11.1) | 8 (5.6) | 15 (10.4) | 3 (2.1) | |
| **Her2** | | | | | 0.843 |
| Equivocal | 2 (1.4) | 0 (0.0) | 0 (0.0) | 0 (0.0) | |
| 0 | 61 (42.4) | 20 (13.9) | 20 (13.9) | 6 (4.2) | |
| 1+ | 3 (2.1) | 1 (0.7) | 1 (0.7) | 0 (0.0) | |
| 2+ | 3 (2.1) | 1 (0.7) | 2 (1.4) | 1 (0.7) | |
| 3+ | 18 (12.5) | 2 (1.4) | 2 (1.4) | 1 (0.7) | |
| **Stage** | | | | | 0.391 |
| I | 5 (3.5) | 2 (1.4) | 1 (0.7) | 2 (1.4) | |
| IIA | 34 (23.6) | 7 (4.9) | 10 (6.9) | 5 (3.5) | |
| IIB | 31 (21.5) | 9 (6.2) | 7 (4.9) | 1 (0.7) | |
| IIIA | 8 (5.6) | 2 (1.4) | 5 (3.5) | 0 (0.0) | |
| IIIB | 9 (6.2) | 4 (2.8) | 2 (1.4) | 0 (0.0) | |
| **Grade** | | | | | 0.273 |
| Missing | 51 (35.4) | 9 (6.2) | 12 (8.3) | 3 (2.1) | |
| 1 | 15 (10.4) | 9 (6.2) | 6 (4.2) | 2 (1.4) | |
| 2 | 21 (14.6) | 5 (3.5) | 7 (4.9) | 3 (2.1) | |
| 3 | 0 (0.0) | 1 (0.7) | 0 (0.0) | 0 (0.0) | |
| **Histological type** | | | | | 0.247 |
| IDC | 37 (25.7) | 15 (10.4) | 13 (9.0) | 5 (3.5) | |
| ILC | 50 (34.7) | 9 (6.2) | 12 (8.3) | 3 (2.1) | |

#### Supplementary Table 2_Tropee_2019

| | SMARCD3 score | | | | |
|---|---|---|---|---|---|
| | - | + | ++ | +++ | p-value |
| **Age (years)** | | | | | 0.154922539 |
| < 45 | 6 (2.9) | 2 (1.0) | 11 (5.3) | 2 (1.0) | |
| ≥ 45 | 49 (23.7) | 37 (17.9) | 58 (28.0) | 42 (20.3) | |
| **PR** | | | | | **0.018490755** |
| Positive | 20 (9.7) | 14 (6.8) | 12 (5.8) | 7 (3.4) | |
| Negative | 35 (16.9) | 25 (12.1) | 57 (27.5) | 37 (17.9) | |
| **T-stage** | | | | | **0.006496752** |
| pT1 | 7 (3.4) | 5 (2.4) | 19 (9.2) | 9 (4.3) | |
| pT2 | 29 (14.0) | 24 (11.6) | 30 (14.5) | 11 (5.3) | |
| pT3 | 13 (6.3) | 9 (4.3) | 17 (8.2) | 22 (10.6) | |
| pT4 | 6 (2.9) | 1 (0.5) | 3 (1.4) | 2 (1.0) | |
| **N-stage** | | | | | 0.817091454 |
| pN1 | 35 (16.9) | 21 (10.1) | 44 (21.3) | 28 (13.5) | |
| pN2 | 15 (7.2) | 13 (6.3) | 18 (8.7) | 9 (4.3) | |
| pN3 | 5 (2.4) | 5 (2.4) | 7 (3.4) | 7 (3.4) | |
| **Histological type** | | | | | 0.784607696 |
| Lobular | 8 (3.9) | 3 (1.4) | 9 (4.3) | 13 (6.3) | |
| NST | 35 (16.9) | 27 (13.0) | 46 (22.2) | 19 (9.2) | |
| Other | 12 (5.8) | 9 (4.3) | 14 (6.8) | 12 (5.8) | |
| **Grade** | | | | | 0.457271364 |
| 1 | 6 (2.9) | 4 (1.9) | 8 (3.9) | 3 (1.4) | |
| 2 | 29 (14.0) | 27 (13.0) | 44 (21.3) | 32 (15.5) | |
| 3 | 20 (9.7) | 8 (3.9) | 17 (8.2) | 9 (4.3) | |
| **Mitotic score** | | | | | 0.417791104 |
| 1 | 27 (13.0) | 26 (12.6) | 44 (21.3) | 31 (15.0) | |
| 2 | 17 (8.2) | 8 (3.9) | 17 (8.2) | 9 (4.3) | |
| 3 | 11 (5.3) | 5 (2.4) | 8 (3.9) | 4 (1.9) | |
| **Multiple** | | | | | 0.217391304 |
| No | 43 (20.8) | 27 (13.0) | 43 (20.8) | 33 (15.9) | |
| Yes | 12 (5.8) | 12 (5.8) | 26 (12.6) | 11 (5.3) | |

#### Supplementary Table 3_Tropee_2019

| | | n (event) | Single Variable Analysis HR (95% CI) | p-value |
|---|---|---|---|---|
| **Age** | | 229 (48) | 1 (0.98-1.03) | 0.835 |
| ***SMARCD3* expression** | High | 113 (14) | 1 | |
| | Low | 94 (26) | 2.11 (1.10-4.06) | **0.024** |
| | Missing | 22 | | |
| **PR status** | Positive | 168 (30) | 1 | |
| | Negative | 60 (18) | 2.026 (1.1-3.73) | **0.023** |
| | Missing | 1 | | |
| **T-stage** | pT1 | 48 (4) | 1 | |
| | pT2 | 100 (24) | 3.38 (1.17-9.75) | **0.024** |
| | pT3 | 68 (16) | 3.14 (1.05-9.42) | **0.041** |
| | pT4 | 13 (4) | 4.95 (1.23-19.83) | **0.024** |
| **N-stage** | pN1 | 139 (20) | 1 | |
| | pN2 | 64 (19) | 1.83 (0.97-3.45) | 0.06 |
| | pN3 | 26 (9) | 2.47 (1.12-5.44) | **0.025** |
| **Histological type** | Lobular | 38 (9) | 1 | |
| | NST | 140 (27) | 0.69 (0.32-1.46) | 0.329 |
| | Other | 51 (12) | 0.97 (0.41-2.32) | 0.951 |
| **Grade** | 1 | 25 (2) | 1 | |
| | 2 | 143 (30) | 2.99 (0.71-12.51) | 0.134 |
| | 3 | 61 (16) | 3.38 (0.78-14.73) | 0.105 |
| **Radiotherapy** | No | 39 (10) | 1 | |
| | Yes | 187 (38) | 0.84 (0.42-1.68) | 0.616 |
| | Missing | 3 | | |
| **Chemotherapy** | No | 31 (8) | 1 | |
| | Yes | 197 (40) | 0.52 (0.24-1.12) | 0.097 |
| | Missing | 1 | | |
| **Neo-adjuvant** | No | 202 (37) | 1 | |
| | Yes | 26 (11) | 2.55 (1.3-5) | **0.007** |
| | Missing | 1 | | |
| **Endocrine** | No | 11 (6) | 1 | |
| | Yes | 217 (42) | 0.31 (0.13-0.72) | **0.007** |
| | Missing | 1 | | |

Abbreviations: CI, confidence interval; HR, hazard ratio.
p-value derived from Wald statistical test
